## Supplementary material for "*In silico, in vitro* and *in cellulo* models for monitoring SARS-CoV-2 spike/human ACE2 complex, viral entry and cell fusion": Sup data

**SUPPLEMENTARY DATA**

A-

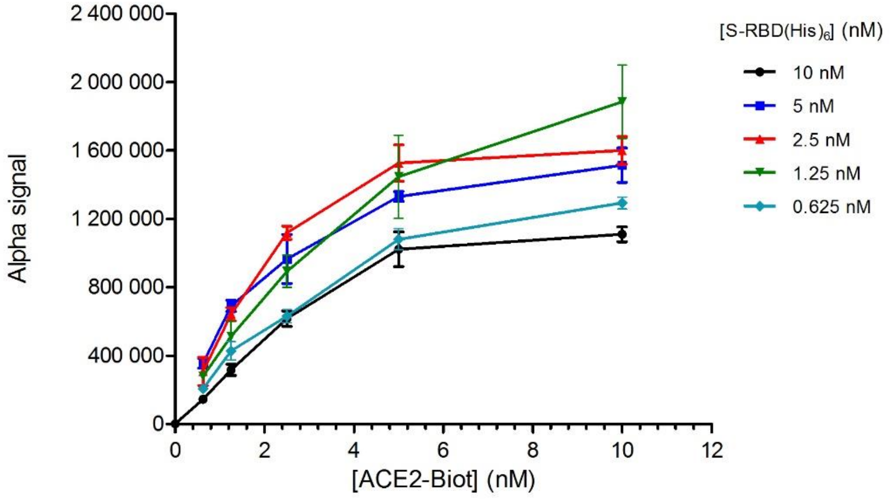


**S1. SARS-CoV-2 S-RBD/ACE2 titration by AlphaLISA.** The AlphaLISA assay development was performed in 96-well ½ aera Alpha plate (Perkin Elmer, Waltham, MA) with a final reaction volume of 40 µL. Cross titration experiments of ACE2-Biot (0.3 to 12 nM) against S-RBD(His)$_6$ (0.625 to 10 nM) were carried out to establish optimal assay concentrations for the binding assay. Data were obtained from at least two independent experiments and are reported as Alpha signal means ±SD.

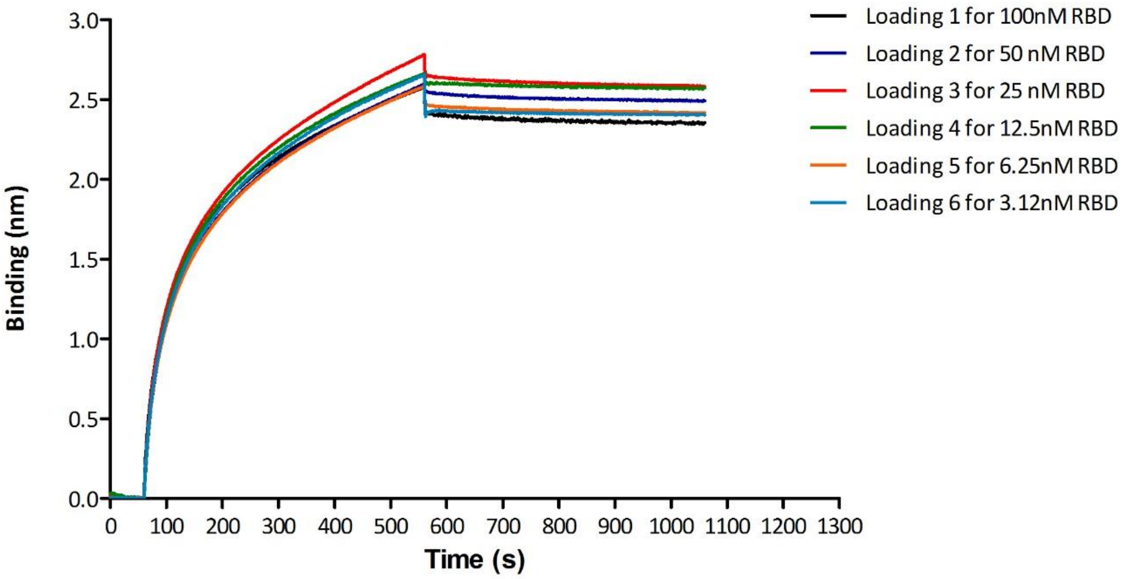


**S2. Loading of 1µM ACE2-biot onto Bio-layer interferometry (BLI) streptavidin biosensors.** Bio-Layer Interferometry (BLI) experiments were performed on a BLItz instrument (ForteBio) to measure the binding of S-RBD(His)$_6$ to ACE2-Biot. Each curve represents the loading of 1µM ACE2-biot onto a streptavidin biosensor for 500s in order to reach a binding value of 2.7 nm for each biosensor. Unbound ACE2 were then removed by a 500s washing step in the reaction buffer. Each Ace2 loaded biosensor was then used for the binding kinetics with RBD-his (from 3.12nM to 100 nM).

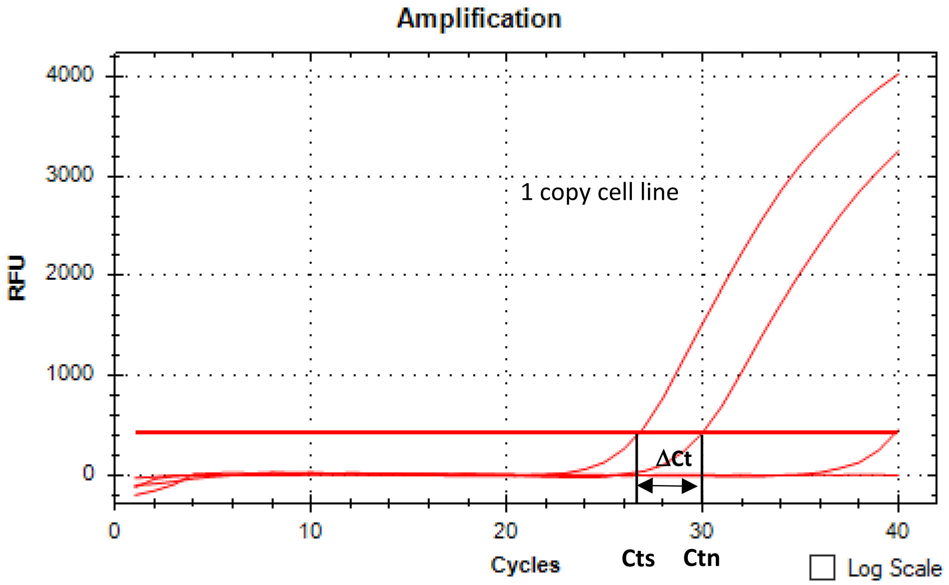


Cts : Ct sample 293T-ACE2
Ctn: Ct normalised cell line 1 copy

| Cell name | Ct | Moy Ct | ΔCT | copies number /cell |
|---|---|---|---|---|
| 393T-ACE2 | 26,78 | 27,071 | -3,173 | 9,021 |
| | 27,35 | | | |
| 293T-NT | N/A | 39,882 | 9,638 | 0,001 |
| | 39,88 | | | |
| H20 | N/A | N/A | N/A | N/A |
| | N/A | | | |
| HEK 2C9 | 30,02 | 30,244 | 0,000 | 1,000 |

Ct sample - Ct standard = ΔCt
Copy number of target gene = 2-ΔCt
1 copy normalised cell line HEK-2C9

**S3. Quantification of the number of *ACE2* copies in HEV293T-ACE2 cells.** HEK293T-ACE2 cell lines have been generated by lentiviral transduction using pHAGE_EF1alnt_ACE2_WT plasmid (kind gift from Bloom's laboratory(10)), see materials and methods section. The efficacy of transduction was assessed using real time PCR ten days post infection. ACE2 provirus DNA from this cell line was quantified by q-PCR using the ΔCt method as compared with 1 copy cell line. DNA from two different cell clones (human 293T and K562 cells), containing a single integrated copy of the provirus, was used as a normalized cell line. HEK293T were cultured in DMEM medium supplemented with gentamycin 5mg/ml and SVF 10%.
